## Supplementary figures and images for "Nanodiamonds enable adaptive-optics enhanced, super-resolution, two-photon excitation microscopy"

### Supplemental Movie 1

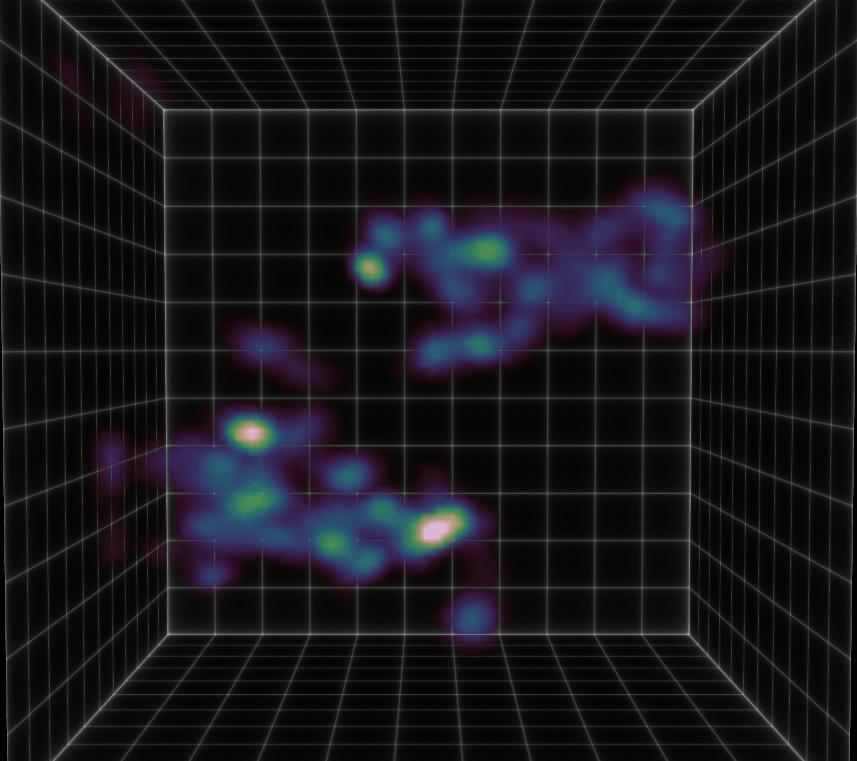
